# Increased metal tolerance and bioaccumulation of zinc and cadmium in *Chlamydomonas reinhardtii* expressing a AtHMA4 C-terminal domain protein

**DOI:** 10.1101/2020.02.13.948307

**Authors:** Aniefon Ibuot, Rachel E. Webster, Lorraine E. Williams, Jon K. Pittman

**Author notes:** Corresponding Author: Dr Jon Pittman, Department of Earth and Environmental Sciences, School of Natural Sciences, The University of Manchester, Michael Smith Building, Oxford Road, Manchester M13 9PT, UK.

## Abstract

The use of microalgal biomass for metal pollutant bioremediation might be improved by genetic engineering to modify the selectivity or capacity of metal biosorption. A plant cadmium (Cd) and zinc (Zn) transporter (AtHMA4) was used as a transgene to increase the ability of *Chlamydomonas reinhardtii* to tolerate 0.2 mM Cd and 0.3 mM Zn exposure. The transgenic cells showed increased accumulation and internalisation of both metals compared to wild type. AtHMA4 was expressed either as the full-length protein or just the C-terminal tail, which is known to have metal binding sites. Similar Cd and Zn tolerance and accumulation was observed with expression of either the full-length protein or C-terminal domain, suggesting that enhanced metal tolerance was mainly due to increased metal binding rather than metal transport. The effectiveness of the transgenic cells was further examined by immobilisation in calcium alginate to generate microalgal beads that could be added to a metal contaminated solution. Immobilisation maintained metal tolerance, while AtHMA4-expressing cells in alginate showed a concentration-dependent increase in metal biosorption that was significantly greater than alginate beads composed of wild type cells. This demonstrates that expressing AtHMA4 full-length or C-terminus has great potential as a strategy for bioremediation using microalgal biomass.

## 1. INTRODUCTION

Metal pollution is a consequence of industrial and mining activities, and causes significant damage to terrestrial and aquatic environments (Boyd, 2010; Kar, Sur, Mandai, Saha, & Kole, 2008). Sustainable and cost-effective biotechnological solutions that would remove toxic metal pollutants are therefore of significant interest (Mani & Kumar, 2014; Ward, 2004). Metals such as cadmium (Cd) and zinc (Zn) are damaging environmental pollutants due to their high toxicity to biota (Järup & Åkesson, 2009; van Straalen, 2002). Microalgae are a potential biomass source for metal pollutant bioremediation within wastewaters or contaminated waters (Mehta & Gaur, 2005; Suresh Kumar, Dahms, Won, Lee, & Shin, 2015). Many natural strains of microalgae are able to tolerate high concentrations of metals and mediate metal biosorption, and a number of studies have examined the capabilities of live and dead microalgal biomass for metal removal from a contaminated water source (Ibuot, Gupta, Ansolia, & Bajhaiya, 2019; Mehta & Gaur, 2005; Monteiro, Castro, & Malcata, 2012; Urrutia, Yañez-Mansilla, & Jeison, 2019; Zeraatkar, Ahmadzadeh, Talebi, Moheimani, & McHenry, 2016). While such studies have demonstrated that the use of microalgal biomass for metal bioremediation is technically feasible, further improvements could be made by enhancing the selectivity and capacity of metal binding and accumulation by microalgae, which could be achieved through genetic engineering (Cheng, Show, Lau, Chang, & Ling, 2019).

Examples of genetic manipulation of microalgae for metal bioremediation are still fairly limited. To date, these include approaches to ectopically express animal metal binding proteins in microalgae, such as a Cd- and mercury (Hg)-binding metallothionein (Cai, Brown, Adhiya, Traina, & Sayre, 1999; He, Siripornadulsil, Sayre, Traina, & Weavers, 2011), manipulation of microalgal amino acid metabolism by expressing histidine or proline biosynthesis genes that enhance metal tolerance (Siripornadulsil, Traina, Verma, & Sayre, 2002; Zheng, Cheng, & Yang, 2013), or enhancement of metal ion reduction such as through the expression of a bacterial mercuric reductase in microalgae to yield a novel method for Hg bioremediation (Huang et al., 2006). An alternative genetic engineering approach aimed to enhance metal accumulation and tolerance in microalgae by over-expression of a metal transporter (Ibuot, Dean, McIntosh, & Pittman, 2017). In this case, a transport protein called CrMTP4 was over-expressed in *Chlamydomonas reinhardtii* and shown to enhance Cd tolerance and total Cd uptake into the microalgal cell. This was proposed to be due to increased transfer and storage of Cd^2+^ into acidic vacuoles (Ibuot et al., 2017). Manipulation of metal transporters for bioremediation is an attractive approach since it may allow both increased metal accumulation into the cell and increased metal tolerance, such as by transfer of a toxic metal out of the cytosol and into an internal compartment.

The P_1B_-type ATPases are a class of metal transporter that are present across all taxa and perform energy-dependent transport of metal ions, including Ag^+^, Cd^2+^, Co^2+^, Cu^2+^, Pb^2+^, Zn^2+^ across cell membranes (Rosenzweig & Argüello, 2012; Williams & Mills, 2005). Members of this family in higher plants are referred to as Heavy Metal ATPases (HMA), and include AtHMA4 from *Arabidopsis thaliana*, which plays a critical role in the transport and homeostasis of Zn and Cd (Hussain et al., 2004; Mills et al., 2005; Mills, Krijger, Baccarini, Hall, & Williams, 2003). AtHMA4 displays *in vivo* localisation at the plasma membrane in the plant (Verret et al., 2004), and predominantly at the plasma membrane with some endoplasmic reticulum (ER) localisation when heterologously expressed in yeast (Baekgaard et al., 2010; Verret et al., 2005). AtHMA4 has a long (473 amino acid) cytosolic C-terminal tail domain that contains a number of di-cysteine residues and a histidine residue repeat (Figure 1a), which has been shown to mediate high-affinity binding of Zn^2+^ and Cd^2+^ (Baekgaard et al., 2010; Ceasar et al., 2020; Lekeux et al., 2018). When only the C-terminal domain of AtHMA4 is expressed in yeast it can provide increased Cd and Zn tolerance (Mills et al., 2010). While previous studies have examined the consequence of expressing the full-length AtHMA4 and/or the C-terminal domain alone in plants such as tobacco and tomato (Kendziorek et al., 2014; Siemianowski et al., 2014; Siemianowski, Mills, Williams, & Antosiewicz, 2011), as well as in yeast, no HMA genes has previously been over-expressed or ectopically expressed in microalgae.

**Figure 1.**
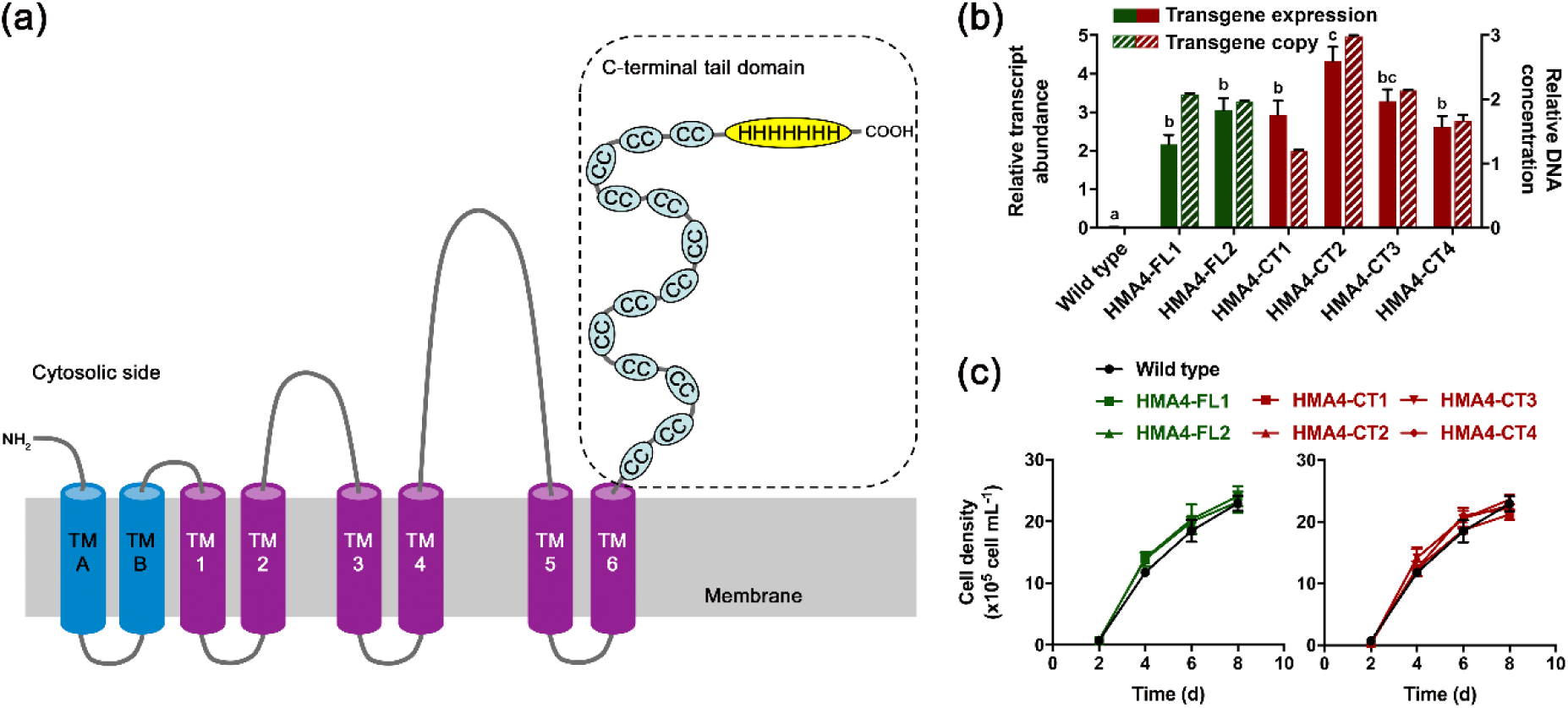
Generation of transgenic *C. reinhardtii* expressing full length (FL) AtHMA4 or the C-terminal domain (CT) of AtHMA4. **(a)** Model of the full length AtHMA4 protein showing the metal binding C-terminal domain possessing 13 paired cysteine residues and a 11 histidine region. **(b)** *AtHMA4* mRNA transcript abundance and transgene copy number in wild type cells in comparison to independent transformant HMA4-FL and HMA-CT lines. qPCR was performed using *CBLP* as a normalization control. **(c)** Cell density of replicate (n=3) empty vector wild type and HMA4-FL and HMA-CT lines over time in standard growth medium.

This study aimed to examine the ability of full-length AtHMA4 expression or AtHMA4 C-terminal domain expression to provide increased tolerance and accumulation of Cd and Zn to the microalga *C. reinhardtii*. This species is still the easiest microalga for genetic manipulation and therefore is an ideal model to examine proof of concept microalgal genetic engineering for bioremediation. For microalgal biomass to be efficiently used for metal bioremediation from polluted water, the biomass should be further modified to facilitate recovery and recycling of the metal from the solution. This can be achieved by processes such as encapsulation and immobilisation of individual cells in a hydrogel matrix (de-Bashan & Bashan, 2010). For example, incorporation of microalgae within alginate can allow the formation of immobilised microalgae-alginate beads and the alginate itself will provide further metal binding capacity (Bayramoğlu & Arica, 2009). Therefore, the efficiency of metal biosorption in the transgenic *C. reinhardtii* following alginate immobilisation was also examined.

## 2. MATERIALS AND METHODS

### 2.1. Microalgae strains and growth conditions

*C. reinhardtii* wild type strain 11/32C was obtained from the Culture Collection of Algae and Protozoa (Oban, UK). *C. reinhardtii* strains expressing full-length or C-terminal domain *AtHMA4* were generated as described in Section 2.2 below. Strains were grown photo-heterotrophically in batch culture in Tris–acetate–phosphate (TAP) medium at pH 7 (Harris, 1989) in 200 mL glass flasks on an orbital shaker rotating at 2 Hz or in 50 mL Nunc flasks, at 25°C under cool-white fluorescent lights (150 μmol m^-2^ s^-1^) with a 16-h:8-h light:dark regime. For metal tolerance and accumulation experiments strains were grown in TAP medium with concentrations of CdCl_2_ (up to 0.2 mM) and ZnSO_4_ (up to 0.3 mM). All cultures were inoculated with the same starting cell density as determined by cell counting to give an initial cell count of 60 × 10^3^ cells mL^-1^.

### 2.2.*C. reinhardtii* nuclear genome transformation and selection

The cDNA encoding the full length AtHMA4 protein or the cDNA encoding the 473 amino acid AtHMA4 C-terminal domain was amplified from previously generated AtHMA4 cDNA plasmids (Mills et al., 2010) by PCR using Phusion High-Fidelity DNA polymerase (New England Biolabs). The primers AtHMA4FL-F (5′-ACT<u>GGATCC</u>CTCTCAACCTTTATCTGAT-3′; BamHI restriction enzyme site underlined) and AtHMA4FL-R (5′-AAA<u>GCGGCCGC</u>ACGTAATGTGAATAGATGGAT-3′; NotI restriction enzyme site underlined) were used to amplify the full length cDNA, and primers AtHMA4CT-F (5′-AAT<u>GGATCC</u>AGGGACTTGTCTGCTTGTGAT-3′; BamHI restriction enzyme site underlined) and AtHMA4CT-R (5′-AAA<u>GCGGCCGC</u>TAATGTGAATAGATGGATGCA-3′; NotI restriction enzyme site underlined) were used to amplify the C-terminal domain cDNA. For all PCR amplification conditions, an annealing temperature of 58°C and 30 amplification cycles were used. Following amplification, the PCR products were BamHI and NotI digested then ligated into the BglII and NotI sites of the Gateway entry plasmid pENTR1A (Thermo Fisher Scientific) for subsequent recombination using an LR Clonase reaction (Thermo Fisher Scientific) into the destination plasmid pH2GW7 (Karimi, De Meyer, & Hilson, 2005) to allow expression of *AtHMA4* in *C. reinhardtii* under control of the constitutive CaMV 35S promoter and hygromycin selection. Correct sequence amplification and ligation was confirmed by DNA sequencing of AtHMA4-FL-pH2GW7 and AtHMA4-CT-pH2GW7 (GATC Biotech).

The pH2GW7 plasmids or empty pH2GW7 plasmid were transformed into *C. reinhardtii* 11/32C using *Agrobacterium tumefaciens* LBA4404 essentially as described previously (Kumar, Misquitta, Reddy, Rao, & Rajam, 2004). Plasmid DNA was transformed into *A. tumefaciens* by freeze-thaw then selection on rifampicin and spectinomycin, and further selection by colony PCR using 35S promoter primers (35SF 5′-GCTCCTACAAATGCCATCA-3′; 35SR 5′-GATAGTGGGATTGTGCGTCA-3′). Selected *A. tumefaciens* strains were propagated then co-cultivated with *C. reinhardtii* cells as described (Kumar et al., 2004). Following co-cultivation, *C. reinhardtii* cells were washed twice with liquid TAP medium containing 500 μg mL^−1^ cefotaxime then transformed lines were selected on TAP agar medium containing 10 μg mL^-1^ hygromycin B and 500 μg mL^−1^ cefotaxime followed by further selection on fresh selection medium. Selected lines were tested by colony PCR using 35S promoter primers and AtHMA4 C-terminal domain primers (CTF 5′-ATGTTGCTGCTGCGAGAGAAGA-3′ and CTR 5′-TCACTTTTTGTTCCCAATCTTTTTCT-3′). Six of the lines named HMA4-FL1, HMA4-FL2, HMA4-CT1, HMA4-CT2, HMA4-CT3 and HMA4-CT4 were studied further and were maintained in selection medium until gene analysis and metal tolerance assays were performed.

Genomic DNA was extracted from the transgenic lines and the control (empty pH2GW7) strain using the cetyl trimethyl ammonium bromide method, exactly as described previously (Bajhaiya, Dean, Zeef, Webster, & Pittman, 2016). RNA was extracted from the strains at day 3 of growth in TAP medium. Cells were harvested by centrifugation at 3000 *g* for 5 min and snap-frozen in liquid N_2_ then RNA extraction was performed using Trizol reagent (Thermo Fisher Scientific), followed by cDNA synthesis using Superscript II reverse transcriptase (Thermo Fisher Scientific) and oligo-dT primer. DNA and RNA concentrations were determined using a NanoDrop UV-Vis spectrophotometer (Thermo Fisher Scientific). Quantitative real-time PCR (qPCR) using a SYBR Green core qPCR kit (Eurogentec) and an ABI Prism 7000 machine (Applied Biosystems) using the SYBR Green detection program was used to determine *AtHMA4* transgene copy number from genomic DNA relative to the single copy *CBLP* gene, also known as *RACK1* (Manuell, Yamaguchi, Haynes, Milligan, & Mayfield, 2005; Schloss, 1990). qPCR was also used to determine *AtHMA4* transcript abundance from cDNA relative to abundance of the constitutive control transcript *CBLP*, which is commonly used as an endogenous reference gene (Castruita et al., 2011). *AtHMA4* was detected using the AtHMA4 C-terminal domain primers described above and *CBLP* was detected using primers CBLPF (5′-CTTCTCGCCCATGACCAC-3′) and CBLPR (5′-CCCACCAGGTTGTTCTTCAG-3′). Reactions were run in triplicate and PCR efficiencies were determined using the comparative threshold cycle method (Schmittgen & Livak, 2008) using LinRegPCR (Ruijter et al., 2009). Melting curves were produced to ensure that single products were amplified. Relative amplification efficiency obtained by qPCR varied between 98-99% and the obtained R^2^ value of all the qPCRs were >0.98.

### 2.3. Microalgae analysis

At regular intervals over 8 d, cultures were grown in TAP medium with or without added Cd or Zn and sampled to determine cell density by cell counting using a Nexcelom Cellometer T4 (Nexcelom Biosciences) or total chlorophyll (chlorophyll a+b) concentration as described previously (Osundeko, Davies, & Pittman, 2013). Internal (sub-cellular) metal content in microalgae after 8 d growth in metal-containing medium was performed following EDTA washing to remove external (cell wall) bound metals by inductively coupled plasma atomic emission spectroscopy (ICP-AES) essentially as described previously (Webster, Dean, & Pittman, 2011). Cells were collected by centrifugation (3000 *g* for 10 min) then resuspended in 10 mL of 10 mM EDTA, incubated for 5 min, then re-centrifuged and washed with 15 mL Milli-Q (Millipore) water. Cell pellets were dried at 60°C for 24 h and then digested in 0.5 mL of ultrapure concentrated nitric acid at 70°C for 2 h. Samples were diluted in Milli-Q water to 2% (v/v) concentration of acid and analysed using a Perkin-Elmer Optima 5300. All samples were calibrated using a matrix-matched serial dilution of Specpure multi-element plasma standard solution 4 (Alfa Aesar) set by linear regression.

### 2.4. Alginate immobilisation and metal exposure

*C. reinhardtii* cells (control and transgenic lines) were cultivated in TAP medium for 6 d until mid to late exponential phase was reached and a cell density of approximately 20 × 10^5^ cells mL^-1^. Cells were further concentrated 5-fold by centrifuging 50 mL of cells at 3000 *g* for 5 min then resuspending the cell pellet in 10 mL of fresh TAP medium. The algal suspension was mixed into an equal volume of 3% (w/v) sodium alginate (Sigma, Product number 180947) solution then a syringe was used to generate 3 – 4 mm diameter algal-alginate beads by adding drops of the sodium alginate-algal mixture into a 2% (w/v) CaCl_2_ solution to generate a solidified calcium alginate. The calcium alginate-algal beads were left in the CaCl_2_ solution for 30 min to allow the beads to harden before they were rinsed in cold deionised water prior to use.

For metal exposure experiments, ten calcium alginate-algal beads were added to 50 mL TAP medium containing CdCl_2_ (0.2 mM, 0.5 mM or 1 mM) and ZnSO_4_ (0.5 mM, 1 mM or 1.5 mM) and incubated for 24 h. For comparison, free-swimming cells of an equal cell density equivalent to the ten beads were exposed to the same metal conditions. Following exposure, the calcium alginate-algal beads or the free-swimming algal cell pellet, collected following centrifugation at 3000 *g* for 5 min, were dried at 60°C for 24 h then digested in nitric acid prior to analysis by ICP-AES, as described above in Section 2.3. For cell growth determination, alginate immobilised cells or free-swimming cells were incubated in 0.2 mM Cd or 0.5 mM Zn media for 3 d. Cell counts were measured on each day. To determine cell counts from alginate beads, beads were incubated in 0.5 M Na-citrate for 30 min to dissolve the hydrogel and release algal cells, which were collected by centrifugation.

### 2.5 Statistical analysis

Data points are presented as mean ± standard error of the mean (SEM). Statistical analysis was performed using GraphPad Prism v.6 by one-way or two-way analysis of variance, as appropriate and using Tukey post-hoc test. Statistically significant difference was considered at *p* < 0.05.

## 3. RESULTS

### 3.1. Ectopic expression of full-length AtHMA4 and AtHMA4 C-terminal tail in *C. reinhardtii*

Two cDNA constructs encoding either full-length AtHMA4 (HMA4-FL) or the C-terminal tail domain of AtHMA4 (HMA-CT) were transformed into *C. reinhardtii* and colonies were selected on the basis of hygromycin tolerance. Ultimately, six stable transgenic lines, including two HMA4-FL lines and four HMA-CT lines were identified that expressed *AtHMA4* transcript sequence in comparison to the absence of any *AtHMA4* expression in the empty-plasmid (wild type) control strain (Figure 1b). Most of the lines had multi-copy insertion of the transgene, either as two copies (HMA4-FL1, HMA4-FL2, HMA4-CT3, HMA4-CT4) or three copies (HMA4-CT2) with just one single-copy insertion (HMA4-CT1), but transgene transcript abundance was equivalent between all lines apart from HMA4-CT2 where expression was significantly higher (Figure 1b). These six lines were taken forward for phenotypic analysis. The expression of the AtHMA4 proteins had no significant influence on *C. reinhardtii* morphology or cell growth under normal non-stressed cultivation conditions. In particular, there was no significant difference in cell density over time of the HMA4-FL and HMA4-CT lines compared to wild type when cells were grown in liquid TAP medium (Figure 1c).

Both HMA4-FL lines expressing the entire AtHMA4 Cd^2+^, Zn^2+^ transporter including the C-terminal tail and all four HMA4-CT lines expressing only the C-terminal tail of AtHMA4 displayed increased Cd tolerance (Figure 2). Cell growth of wild type *C. reinhardtii* was significantly inhibited by the addition of 0.2 mM Cd, while growth of the HMA4-FL lines and the HMA4-CT lines remained strong in the presence of Cd. For all HMA4-FL and HMA4-CT lines cell density was significantly higher than wild type. There was no difference in cell density for the transgenic lines in the absence and presence of Cd, with the exception of the HMA4-CT2 line, which although still higher than wild type had a slight reduction in cell density when grown in Cd medium (Figure 2b). The presence of Cd caused chlorosis and a significant reduction in total chlorophyll content in the wild type after 8 d growth (Figure 2c). In contrast, all HMA4-FL and HMA4-CT lines had significantly higher chlorophyll content compared to wild type in the Cd growth conditions.

**Figure 2.**
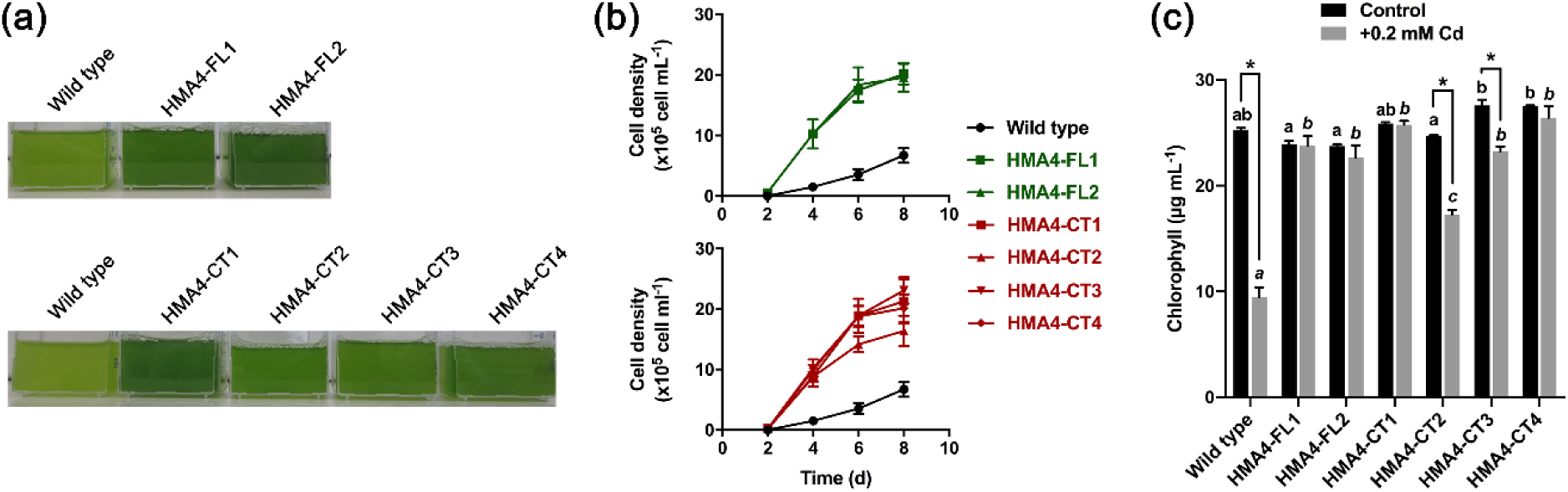
Cd tolerance of HMA4-FL and HMA-CT lines. **(a)** Culture phenotypes of empty vector wild type *C. reinhardtii* and AtHMA4 full length (FL) and C-terminal domain (CT) lines after 8 d growth in TAP medium containing 0.2 mM Cd. A representative experiment is shown. **(b)** Cell density as determined by cell count measurement over time in empty vector wild type and HMA-FL and HMA-CT lines in TAP medium with 0.2 mM Cd addition. **(c)** Total chlorophyll yield in empty vector wild type and HMA-FL and HMA-CT lines in TAP medium with 0.2 mM Cd addition after 8 d growth. Chlorophyll values are shown in comparison to control treatment without Cd addition. Data points are means (±SE) of 3 independent biological replicates. Bars indicated by different lower case letters show significant difference (*P* < 0.05) within treatments (italics letters for Cd treatment; non-italics letters for control treatment) between wild type and transgenic lines. Bars indicated by an asterisk show significant difference between control and Cd treatments (*P* < 0.05).

The HMA4-FL and HMA4-CT lines also showed a consistent increased Zn tolerance compared to wild type control (Figure 3). While the Zn treatment used here (0.3 mM Zn addition) significantly reduced the cell density of all lines in comparison to growth in control growth medium without Zn, all of the HMA4-FL and HMA4-CT lines were more tolerant to the Zn exposure on the basis of cell density compared to the wild type (Figure 3b). Likewise, the Zn addition induced a reduction in total chlorophyll in all lines, but chlorophyll content was significantly higher in all transgenic microalgae lines compared to wild type in the presence of Zn (Figure 3c). Together these data indicate that the expression of both full length AtHMA4 and the just the C-terminal tail domain can provide substantial tolerance to Cd and Zn toxicity for *C. reinhardtii*.

**Figure 3.**
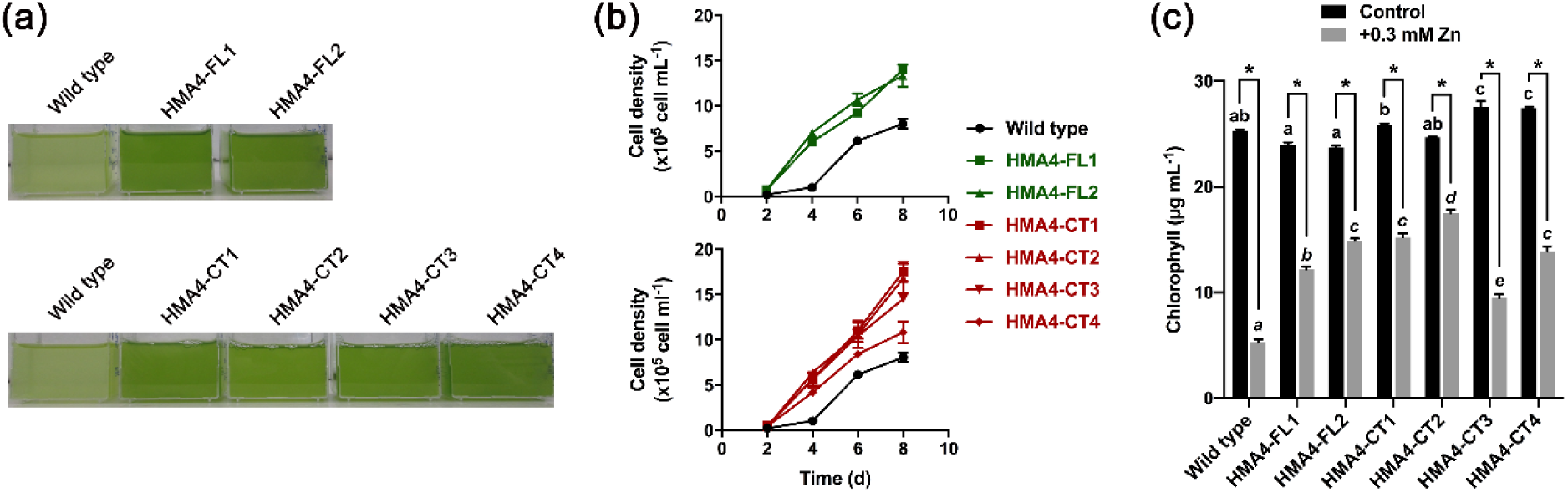
Zn tolerance of HMA4-FL and HMA-CT lines. **(a)** Culture phenotypes of empty vector wild type *C. reinhardtii* and AtHMA4 full length (FL) and C-terminal domain (CT) lines after 8 d growth in TAP medium containing 0.3 mM Zn. A representative experiment is shown. **(b)** Cell density as determined by cell count measurement over time in empty vector wild type and HMA-FL and HMA-CT lines in TAP medium with 0.3 mM Zn addition. **(c)** Total chlorophyll yield in empty vector wild type and HMA-FL and HMA-CT lines in TAP medium with 0.3 mM Zn addition after 8 d growth. Chlorophyll values are shown in comparison to control treatment without Zn addition. Data points are means (±SE) of 3 independent biological replicates. Bars indicated by different lower case letters show significant difference (*P* < 0.05) within treatments (italics letters for Zn treatment; non-italics letters for control treatment) between wild type and transgenic lines. Bars indicated by an asterisk show significant difference between control and Zn treatments (*P* < 0.05).

### 3.2. Cd and Zn uptake into *C. reinhardtii*

Cd and Zn concentration in algal cell biomass was determined by ICP-AES measurement in order to determine whether the expression of full-length AtHMA4 or the AtHMA4 C-terminal tail can give rise to increased metal content within the cell. Prior to measurement, cells were grown in liquid media with added Cd or Zn then washed with the metal chelator EDTA to remove cell wall bound metals so that only internalised metals were measured. All of the transgenic lines except HMA4-CT2 showed a significant, approximately 2-fold increase in Cd content in comparison to wild type control (Figure 4a), while all of the transgenic *C. reinhardtii* lines except HMA4-CT4 showed an approximately 2- to 3-fold significant increase in Zn content within the cells (Figure 4b). This suggests that the Cd and Zn tolerance of the AtHMA4 transgenic lines was potentially due to internal sequestration of metal rather than efflux from the cell.

**Figure 4.**
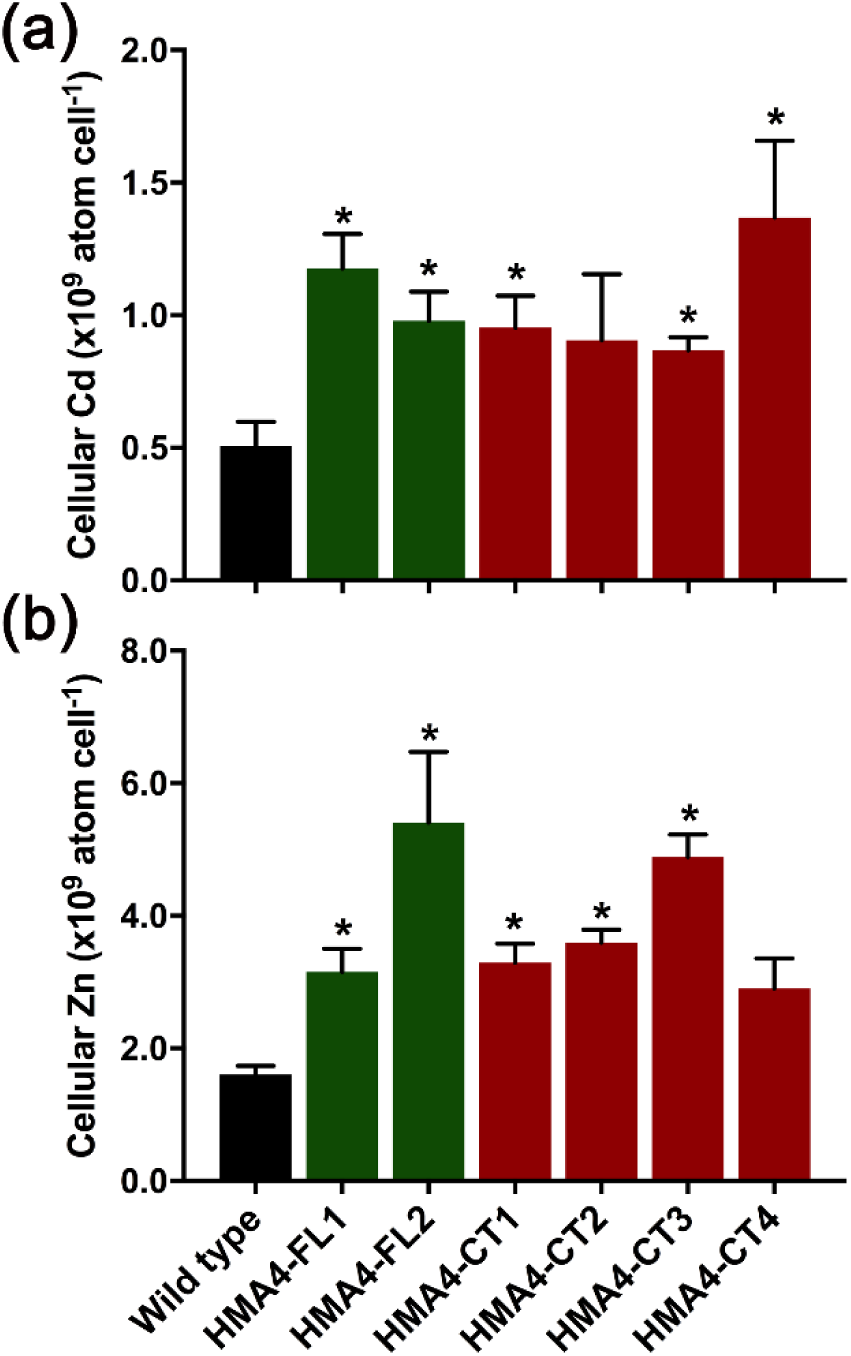
Cd and Zn accumulation of HMA4-FL and HMA-CT lines. **(a)** Cellular Cd content in EDTA-washed cells following growth in 0.2 mM Cd medium after 8 d. **(b)** Cellular Zn content in EDTA-washed cells following growth in 0.3 mM Zn medium after 8 d. Data points are means (±SE) of 3 independent biological replicates. Bars indicated by an asterisk show significant difference (*P* < 0.05) with the wild type strain.

### 3.3 Alginate immobilisation of *C. reinhardtii*

In order to evaluate the viability of the genetically engineered cells to accumulate metals when in an immobilised form, the cells were mixed with sodium alginate then gelled with use of calcium to produce calcium alginate microalgal beads. A comparison is shown between untreated (free-swimming) and alginate immobilised cells of wild type *C. reinhardtii*, the full-length AtHMA4 expression line, HMA4-FL2, and the C-terminal domain line, HMA4-CT1. Immobilisation slightly reduced cell growth rate in all cases although this reduction was only significant for the wild type cells grown under no metal exposure conditions (Figure 5). The enhanced tolerance to Cd and Zn by the HMA4-FL2 and HMA4-CT1 cells in comparison to wild type was maintained following the immobilisation process as growth rates of the transgenic cells in the presence of alginate were significantly higher than the immobilised wild type cells when incubated in either metal.

**Figure 5.**
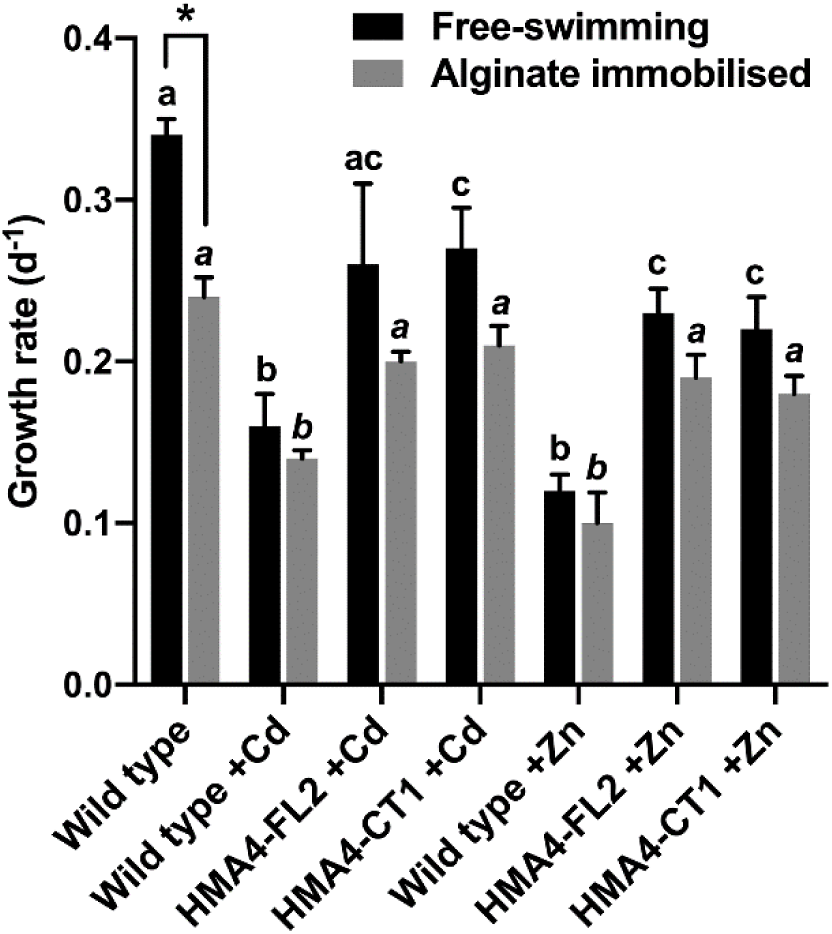
Alginate immobilisation of *C. reinhardtii* cells. Growth rate of free-swimming cells and alginate-immobilised cells, either empty vector wild type, HMA4-FL2 line or HMA4-CT1 line, determined after 3 d of growth in standard TAP medium with metal addition, or in medium with 0.2 mM Cd or 0.5 mM Zn addition. Data points are means (±SE) of 3 independent biological replicates. Bars indicated by different lower case letters show significant difference (*P* < 0.05) within treatments (italics letters for alginate immobilised cells; non-italics letters for free-swimming cells) between wild type and transgenic lines. Bars indicated by an asterisk show significant difference between free-swimming and alginate-immobilised cells (*P* < 0.05).

Calcium alginate alone is able to bind metals including Cd and Zn from solution (Jodra & Mijangos, 2001), therefore total metal biosorption of microalgae-calcium alginate beads was determined alongside total metal biosorption of free-swimming algal biomass without EDTA washing. Metal biosorption of the alginate microalgal beads was initially examined over 3 d following addition of the beads into metal containing medium at a range of Cd and Zn concentrations. Maximal biosorption of both Cd and Zn to the beads was seen within 24 h while biosorption values were highly variable at earlier time points. Furthermore, the free-swimming cells exhibited severe chlorosis and cell death at the higher Cd and Zn concentrations (1 mM Cd, 1.5 mM Zn) after 24 h, therefore all measurements were performed at the 24 h time period. Due to the additional metal binding capacity of the calcium alginate, all immobilised cells displayed an approximately 5 – 10 times higher Cd and Zn metal biosorption in comparison to the free-swimming cells (Figure 6). There was a concentration-dependent increase in Cd and Zn biosorption for both the free-swimming cells and the immobilised cells, except for the free-swimming HMA4-FL2 and HMA4-CT1 cells exposed to 1 mM Cd where there was no further increase in Cd biosorption compared to the 0.5 mM Cd treatment. Furthermore, the immobilised transgenic cells all displayed a significant increase in metal biosorption in comparison to the immobilised wild type cells for each metal treatment, and similar results were observed for the HMA4-FL2 and HMA4-CT1 cells (Figure 6).

**Figure 6.**
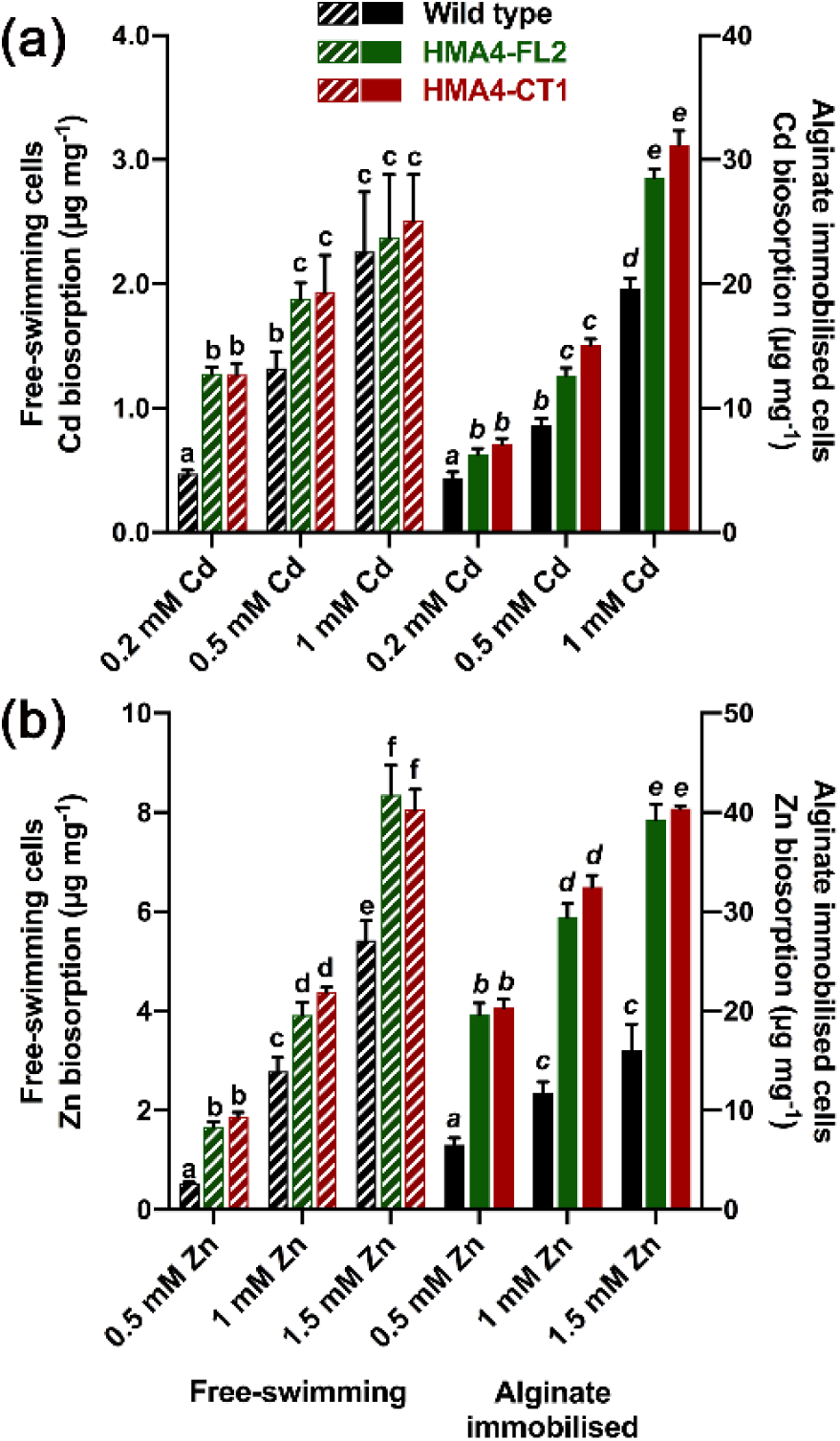
Total metal content of alginate immobilised *C. reinhardtii* cells in comparison to free-swimming cells. A dense sample of late exponential phase free-swimming cells or alginate beads containing cells, either empty vector wild type, HMA4-FL2 line or HMA4-CT1 line, were added to TAP medium containing concentrations of Cd **(a)** or Zn **(b)**. Total Cd or Zn content of harvested cells (free-swimming cells) or beads (alginate immobilised cells) was determined following 24 h incubation in metal medium. Data points are means (±SE) of 3 independent biological replicates. Bars indicated by different lower case letters show significant difference (*P* < 0.05) within treatments (italics letters for alginate immobilised cells; non-italics letters for free-swimming cells) between wild type and transgenic lines.

## 4. DISCUSSION

The genetic manipulation of microalgae in order to increase toxic metal tolerance and accumulation for possible bioremediation applications requires the selection and successful expression of a suitable candidate gene. The *A. thaliana* heavy metal transporter AtHMA4 has been previously considered as an attractive gene for genetically engineering plants for bioremediation (Siemianowski et al., 2011; Verret et al., 2004), and although this gene has also been used to increase metal tolerance in yeast (Mills et al., 2005; Mills et al., 2010), it has never been expressed in microalgae. Here we have demonstrated that the ectopic expression in *C. reinhardtii* of either the full-length AtHMA4 or the C-terminal tail of AtHMA4 was able to provide clear tolerance to Cd and Zn, which was consistent with the known *in vivo* substrate specificity of AtHMA4 (Hussain et al., 2004; Mills et al., 2005; Mills et al., 2003). The degree of Cd and Zn tolerance was generally equivalent regardless of whether the full-length AtHMA4 or C-terminal AtHMA4 construct was expressed. This is different to phenotypes previously seen in yeast where expression of the C-terminal tail by itself gave significantly greater Cd and Zn tolerance than in yeast expressing full-length AtHMA4 (Baekgaard et al., 2010; Mills et al., 2005; Mills et al., 2010). Likewise, enhanced Cd and Zn tolerance in yeast was also seen when the C-terminal tail of AhHMA4 from *Arabidopsis halleri* was expressed (Courbot et al., 2007). The key difference between the transgenic *C. reinhardtii* and the transgenic yeast phenotypes is likely due to the functioning of full-length AtHMA4 in these cells in comparison to the C-terminus.

It has been proposed that the C-terminal domain alone, which has no transport function, can provide metal tolerance by acting as a Cd^2+^ and Zn^2+^ chelator and therefore lower the concentration of free cation within the cytosol despite an increase in total metal content (Baekgaard et al., 2010; Mills et al., 2010). The C-terminal domain exhibits high-affinity Zn^2+^ binding (*K*_D_ < 10 nM) and a stoichiometry of 10 – 11 Zn(II) atoms, predominantly due to the di-cysteine residues (Baekgaard et al., 2010; Lekeux et al., 2018), while high-affinity Cd^2+^ binding (*K*_D_ < 126 nM) to the C-terminal domain has a stoichiometry of approximately 4 Cd(II) atoms but only partial association with the di-cysteine residues (Ceasar et al., 2020). In contrast, full-length AtHMA4 is an ATP-dependent active cation transporter that will drive Cd^2+^ and Zn^2+^ transport across a membrane against an electrochemical gradient. In *A. thaliana* AtHMA4 is located at the plasma membrane within root vascular cells allowing the efflux of metals out of the xylem parenchyma cells for subsequent translocation into the plant shoots (Hussain et al., 2004; Verret et al., 2004). Likewise, the partial plasma membrane localisation of AtHMA4 in wild type yeast and reduced Cd and Zn content indicates that the transporter is mediating metal efflux out of the yeast cell (Mills et al., 2005; Verret et al., 2005). This cellular efflux of metals has been proposed to explain the moderate gain of tolerance by yeast expressing full-length AtHMA4. In contrast, AtHMA4 expression in *C. reinhardtii* increased total Cd and Zn content suggesting that the protein was not localised at the plasma membrane in the algal cell and therefore unable to perform metal efflux or that metal efflux activity was inhibited. This is similar to what was seen when full-length AtHMA4 was expressed in a *zrc1 cot1* mutant yeast strain causing an increase in total Cd content (Baekgaard et al., 2010). Therefore, when expressed in *C. reinhardtii*, AtHMA4 may be localised at an internal compartment thereby allowing detoxification by metal sequestration. Alternatively, the normal transport function could be inhibited and so the protein allows metal tolerance by a different mechanism. The identical phenotypes of the *C. reinhardtii* HMA4-FL and HMA-CT lines suggests that both forms of the AtHMA4 protein provide metal tolerance by metal binding to the C-terminal tail rather than any metal transport, allowing the cells to tolerate increased levels by chelating the cellular free metal ions.

Previous *C. reinhardtii* genetic engineering studies have also enhanced metal tolerance due to the introduction of transgenes that increase intracellular metal binding processes. These include expression of a pyrroline-5-carboxylate synthetase (P5CS) gene that increases phytochelatin synthesis and thus Cd sequestration due to higher proline content (Siripornadulsil et al., 2002). In this study the transgenic P5CS cells could tolerate 0.1 mM Cd and could bind 4-fold more Cd than wild type. Another study showed increased tolerance to 40 μM Cd and a modest increase in Cd removal from the medium due to expression of a chicken Cd-binding metallothionein (Cai et al., 1999). Collectively our work here and these previous studies clearly show that manipulation of intracellular proteins with metal binding characteristics can play a key role in enhancing metal tolerance and accumulation within microalgae.

To allow microalgal biomass to be used for wastewater or contaminated water bioremediation on a commercial scale, an immobilised system is particularly attractive because it minimises the challenge of recovering the metal-containing biomass from solution, it prevents the microalgae itself becoming a pollutant, and it eases the recycling of both metal and biosorbant (de-Bashan & Bashan, 2010; Mallick, 2002). A natural polymer such as alginate, typically derived from brown algae, is one of the most commonly used materials for microalgal encapsulation, and is attractive due to being non-toxic and having high metal-binding capacity (Jodra & Mijangos, 2001). Many studies have evaluated and described the metal removal characteristics of alginate beads derived from various microalgae species including *C. reinhardtii, Chlorella vulgaris, Dunaliella salina* and *Scenedesmus quadricauda* (Bayramoğlu & Arica, 2009; Bayramoğlu, Tuzun, Celik, Yilmaz, & Arica, 2006; Mehta & Gaur, 2001; Moreno-Garrido, Campana, Lubián, & Blasco, 2005). All of these analyses used natural strains whereas here we examined the alginate immobilisation of transgenic microalgae. While the development of a genetically modified microalga for metal bioremediation may be primarily of value as a proof of concept for enhancing specific metal accumulation characteristics, if a transgenic strain was to be exploited it would certainly require containment such as by alginate immobilisation within a bioreactor. Therefore the demonstration here that the significant metal accumulation characteristics of modified HMA4-FL and HMA4-CT *C. reinhardtii* strains are not inhibited by alginate immobilisation is an important validation.

## ACKNOWLEDGEMENTS

This work was financially supported by a Government of Nigeria TETFUND PhD studentship awarded to A.I. and supported in part by The Leverhulme Trust grant (grant number F/00 120/BG) awarded to J.K.P. We thank Paul Lythgoe (Manchester Analytical Geochemical Unit, Department of Earth and Environmental Sciences, University of Manchester) for ICP-AES analysis. We thank Jessica Needham, Zizhen Wan and Silei Lu for assistance in performing experiments with alginate immobilised microalgae.

## AUTHOR CONTRIBUTIONS

All authors contributed towards the design of experiments and analysis of experimental data, A.I., R.E.W., and J.K.P. generated experimental data, J.K.P. wrote the manuscript draft, all authors read, edited and approved the final manuscript.

